# Gene mapping of nine agronomic traits and genome assembly by resequencing a foxtail millet RIL population

**DOI:** 10.1101/069625

**Authors:** Xuemei Ni, Qiuju Xia, Houbao Zhang, Shu Cheng, Hui Li, Guangyu Fan, Tao Guo, Ping Huang, Haitao Xiang, Qingchun Chen, Ning Li, Hongfeng Zou, Xuemei Cai, Xuejing Lei, Xiaoming Wang, Chengshu Zhou, Zhihai Zhao, Gengyun Zhang, Zhiwu Quan

**Affiliations:** BGI-Shenzhen, Shenzhen 518083, China.; Institute of millet, Zhangjiakou Academy of Agricultural Science, Zhangjiakou 075000, China.; State Key Laboratory of Agricultural Genomics, BGI-Shenzhen, Shenzhen 518083, China.; Key Lab of Genomics, Chinese Ministry of Agriculture, BGI-Shenzhen, Shenzhen 518083, China.; Guangdong Province key laboratory of crop germplasm research and application, BGI-Shenzhen, Shenzhen 518083, China.; Shenzhen engineering laboratory of molecular design breeding, BGI-Shenzhen, Shenzhen 518083, China.

## Abstract

Foxtail millet (*Setaria italica*) provides food and fodder in semi-arid regions and infertile land. Resequencing of 184 foxtail millet recombinant inbred lines (RILs) was carried out to aid essential research on foxtail millet improvement. Bin map were constructed based on the RILs’ recombination data. By anchoring some unseated scaffolds and filling gaps, we update two original millet reference genomes Zhanggu and Yugu to produce second editions. Gene mapping of nine agronomic traits were done based on this RIL population. The genome resequencing and QTL mapping provided important tools for foxtail millet research and breeding. Resequencing of the RILs could also provide an effective way for high quantity genome assembly and gene identification.

## INTRODUCTION

Foxtail millet (*Setaria italica*) was an ancient cultivated crop domesticated in China more than 8,700 years ago (ZOHARY and HOPF 2000; BARTON *et al.* 2009). It provided the most important food and forage for the Yellow River valley in ancient China and is still as an essential food source in semi-arid areas (HARLAN 1975; BETTINGER *et al.* 2010). Although foxtail millet is one of the most drought-resistant crops, the low productivity of unimproved foxtail millet has limited its application in agriculture. In the past decade, herbicide resistant, high yield hybrid millet which can dramatically improve crop production and decrease labor cost, indicating that millet has the potential for becoming a high yield crop through the use of genetic tools (DEKKER 2003; SILES *et al.* 2004; AUSTIN 2006).

Millet genome de *novo* sequencing was finished in 2012 (ZHANG *et al.* 2012; BENNETZEN *et al.* 2012), here we carried out resequencing of foxtail millet RIL population and construction of the high resolution bin map. A high accuracy Zhanggu millet genome reference (anchoring 96% scaffold sequence) was constructed based on bin map and Zhanggu draft genome (ZHANG *et al.* 2012). Additionally, this bin map was also used to check the assembly error in Yugu genome reference (BENNETZEN *et al.* 2012), a Yugu genome reference second edition (anchoring 99.7% scaffold sequence) was done after assembly error correction and gap filling. The RIL population was phenotype appraisal for nine agronomic traits in 2010 and 2011, respectively. Gene mapping and QTL analysis was done based on genotype and phenotype date of these RILs, some loci were mapped at 130 Kb zone using 184 RILs, indicating that resequencing of foxtail millet RIL population could provide an effective approach for high quantity genome assembly and gene mapping. The genome reference and SNP markers will become an important tool in millet molecular breeding, the loci related to nine agronomic traits will provide pivotal information to millet breeders.

## MATERIALS AND METHODS

No specific permissions were required for the described field studies. The location i s not privately owned or protected, and the field studies did not involve endangered or protected species.

### Sampling and RIL construction

Zhanggu was selected as the male parent line and A2 was selected as the male sterile line. F1 was constructed from a cross between Zhanggu and A2, RILs was developed by using single seed descent strategy. Segregation population was grown three generation per year in New Village, Jiyang Town, Sanya City, Hainan province (Coordinates: 109°35’E/18°17’N) (November-January; January-April) and Erliban Village, Shalingzi Town, Xuanhua County, Zhangjiakou City, Hainan province (Coordinates: 114°54’E/40°40’N) (May-October).

### Phenotyping

Flag leaf length and width were measured at the maximal values for each flag leaf using a ruler. Plant height was measured the distance between tassel terminal and ground. Additionally, ear height was measured the distance between flag leaf ear and ground. Heading date was recorded as the number of days from sowing to heading.

Sethoxydim resistant data was collected by seeding on Sethoxydim medium (50mg/L). Leaf color was observed and record 15days after sowing, leaf color can divide into yellow and green. Bristle color was observed and record 10days after heading, bristle color can divide into red and green. Anther color was observed and record when flower, anther color can divide into yellow and brown. Tassel hardness was observed and record 30 days after flowering, tassel can divide into stiff and flexible.

### DNA isolation and genome sequencing

Total genomic DNA was extracted from young leaf tissues using the CTAB method (MURRAY and THOMPSON 1980). 500 bp pair-end libraries was constructed under standard protocol provide by Illumina (San Diego, USA). The sequencing was performed using Hiseq 2000 for pair-end 50-cycle sequencing according to the manufacturer’s standard protocol. Low quality reads, reads with adaptor sequences, duplicated reads were filtered and the remained high quality data was used in SNP calling.

### Sequence alignment, genotyping and recombination breakpoint determination

Reads of A2 was mapping to Zhanggu scaffold by using SOAP2 (LI *et al*. 2009) (vision 2.20), SNP calling was conducted by using SAMtools (LI *et al*. 2009) (version 0.1.8) and realSFS (version 0.983). Positions of these SNPs were marked for RIL SNP calling. After mapping the reads of each RIL back to the scaffolds of Zhanggu, SAMtools (LI *et al*. 2009) (version 0.1.8) and realSFS (version 0.983) were used to identify SNPs in each RIL. A sliding window approach was used to evaluate 15 consecutive SNPs for genotype calling and continued the process as the window slide base-by-base (WU *et al*. 2008). The window with a Zhanggu:A2 SNPs ratio of 11:4 or higher was called Zhanggu, 4:11 or lower called A2, SNPs ratio between 11:4 and 4:11 was called heterozygous. The breakpoint was determined at the boundary of the Zhanggu, A2 and heterozygous.

### Bin map and chromosome construction

All SNP data of the 184 RILs were aligned to a matrixs, the minimal interval of two recombination positions was set as 50kb. Adjacent intervals with the same genotype across the 184 RILs were defined as a single recombination bin (WU *et al*. 2008). 3437 recombination bins were serving as 3437 genetic markers, linkage map was constructed using MSTMap (WU *et al*. 2008). New chromosomes were constructed based on bin map.

### Gene mapping and QTL mapping

Phenotype of each RIL and genotype of each bin was collected for gene mapping and QTL analysis. QTL were identified using composite interval mapping performed in the software package MapQTL 5 (VAN OOIJEN and KYAZMA 2004).The likelihood ratio statistic was computed every bin, QTL were called for LOD values higher than 3.0.

### Yugu genome construction and gap filling

Reads of A2 and RILs were mapping to the chromosomes and scaffolds of Yugu (BENNETZEN *et al*. 2012), SNPs between RILs and Yugu were identified using SOAPsnp (LI *et al*. 2009) (Version 1.02). SNP identical to A2 was considered as A2, SNP contrary to A2 was considered as Zhanggu. Bin map was constructed using the same strategy as before, bins with abnormal linkage were move to proper position according to the bin map. Gaps with two flank sequence was mapping to Zhanggu genome, a gap was filled by Zhanggu sequence only when both flank sequence was match to Zhanggu sequence. Zhanggu sequence filled in Yugu gap was shown in lowercase (Fig. 2).

## RESULTS

Two foxtail millet inbred line were selected as the parents: Zhanggu as the male parent line and A2 as the male sterile line. RILs were developed by hybridization between Zhanggu and A2 followed by self-fertilization to F10. Nine agronomic traits were measured for each RIL in 2010 and 2011, respectively.

DNAs were isolated from each RIL’s young leaves using CTAB method (MURRAY and THOMPSON 1980). 500 bp pair-end libraries were constructed according to the standard protocol provide by Illumina (San Diego, USA). The sequencing was performed using Hiseq 2000 for pair-end 50-cycle sequencing according to the manufacturer’s standard protocol. Low quality reads, adaptor sequences and duplicated reads were filtered, the remaining high quality data was used for SNP calling. In total 140 Gb clean data was generated, which given a 2X coverage of each RIL. Genetic variations were detected by comparision between the genome sequence of Zhanggu and resequencing (∼10X) of A2 (ZHANG *et al*. 2012). Reads of A2 were mapped onto Zhanggu scaffolds using SOAP2 (LI *et al*. 2009) (version 2.20), SNP calling was conducted using SAMtools (LI *et al*. 2009) (version 0.1.8) and realSFS (version 0.983). Positions of these SNPs were marked for RIL SNP calling. After mapping the reads of each RIL back to the scaffolds of Zhanggu, SNPs in the marked position were selected for genotyping. A total of 483,414 SNPs were detected, given an average density of 1.2 SNPs per Kb for the RILs (Table 1, Fig. 1).

**Figure 1.**
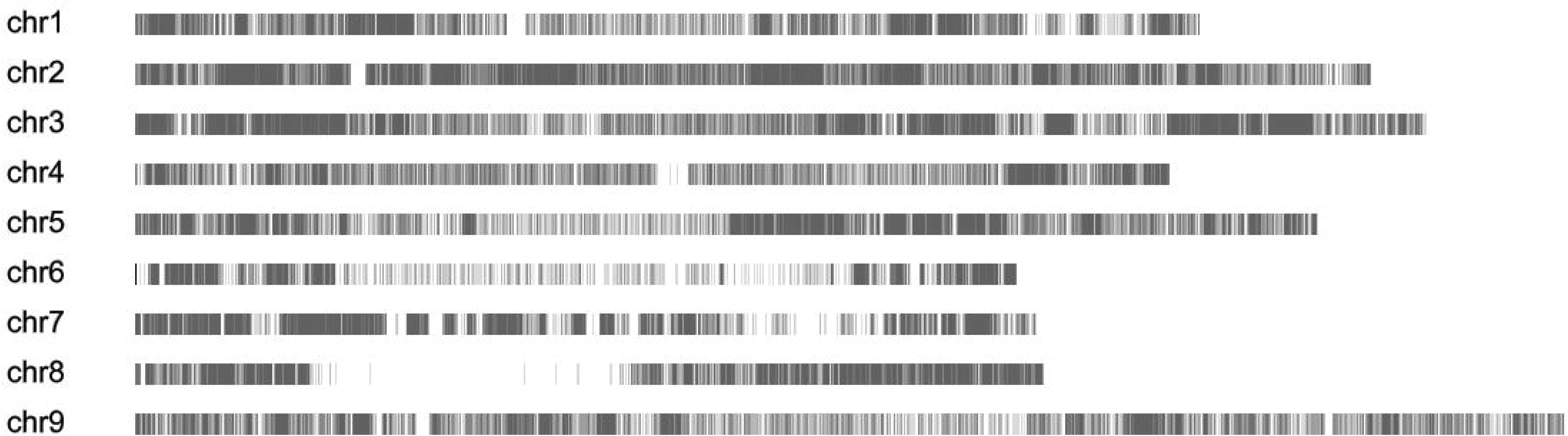
Distribution of 483,414 SNPs between Zhanggu and A2.

**Table 1.** Summary of the bins and SNP distribution in Zhanggu

| Chromosome | Length(bp) | Bin num | Linkage(cM) | SNP num | SNP density(/Kb) |
| --- | --- | --- | --- | --- | --- |
| chr1 | 44603498 | 408 | 167.117 | 42624 | 0.955 |
| chr2 | 51761675 | 491 | 224.647 | 89674 | 1.732 |
| chr3 | 54090027 | 396 | 221.738 | 81236 | 1.502 |
| chr4 | 43349090 | 348 | 168.232 | 27932 | 0.644 |
| chr5 | 49560508 | 480 | 267.52 | 60305 | 1.217 |
| chr6 | 36928436 | 203 | 152.1 | 30946 | 0.838 |
| chr7 | 37743793 | 242 | 168.48 | 56045 | 1.485 |
| chr8 | 38066565 | 283 | 205.431 | 58149 | 1.528 |
| chr9 | 59875680 | 586 | 352.58 | 36503 | 0.609 |
| Total | 415979272 | 3437 | 1927.845 | 483414 | 1.162 |

To avoid potential sources of sequence errors and mapping errors, we chose a sliding window approach to evaluate 15 consecutive SNPs for genotype calling and continued the process as the window slide base-by-base (WU *et al*. 2008). To improve the quality of millet draft genome and conduct genetic analyses, a bin map was constructed based on the recombination data (WU *et al*. 2008). All SNP data of the 184 RILs were aligned to a matrixs, the minimal interval of two recombination positions was set as 10kb. Adjacent intervals with the same genotype across the 184 RILs were defined as a single recombination bin (WU *et al*. 2008). We identify 3437 recombination bins in the 184 RILs, the physical length of the recombination bins ranging from 1 kb to 12Mb, given an average length of 121Kb (Table S1). 3437 single recombination bin were serving as 3437 markers, linkage map was constructed using MSTmap (VAN OOIJEN and KYAZMA 2004). 9 linkage groups, with a total genetic distance of 1927.8 cM (Table 1, Fig 2), were constructed of the foxtail millet genome. The interval between these bins ranging from 0.1 cM to 13.8 cM, average at 0.56 cM. New chromosomes were constructed based on bin map and scaffolds of Zhanggu, and the second edition Zhanggu reference genome was generated (416Mb) after adding unanchored scaffolds (16Mb) (Table 2, Fig. 4). Reads of RILs were mapping to the chromosomes and scaffolds of Yugu to construct bin map, assembly error was revised based on bin map. 3158 gaps was filled by ZhangGu sequence using sequence homology BLAST, the second edition YuGu reference genome was constructed after assembly error correction and gap filling (Fig. 3).

**Figure 2.**
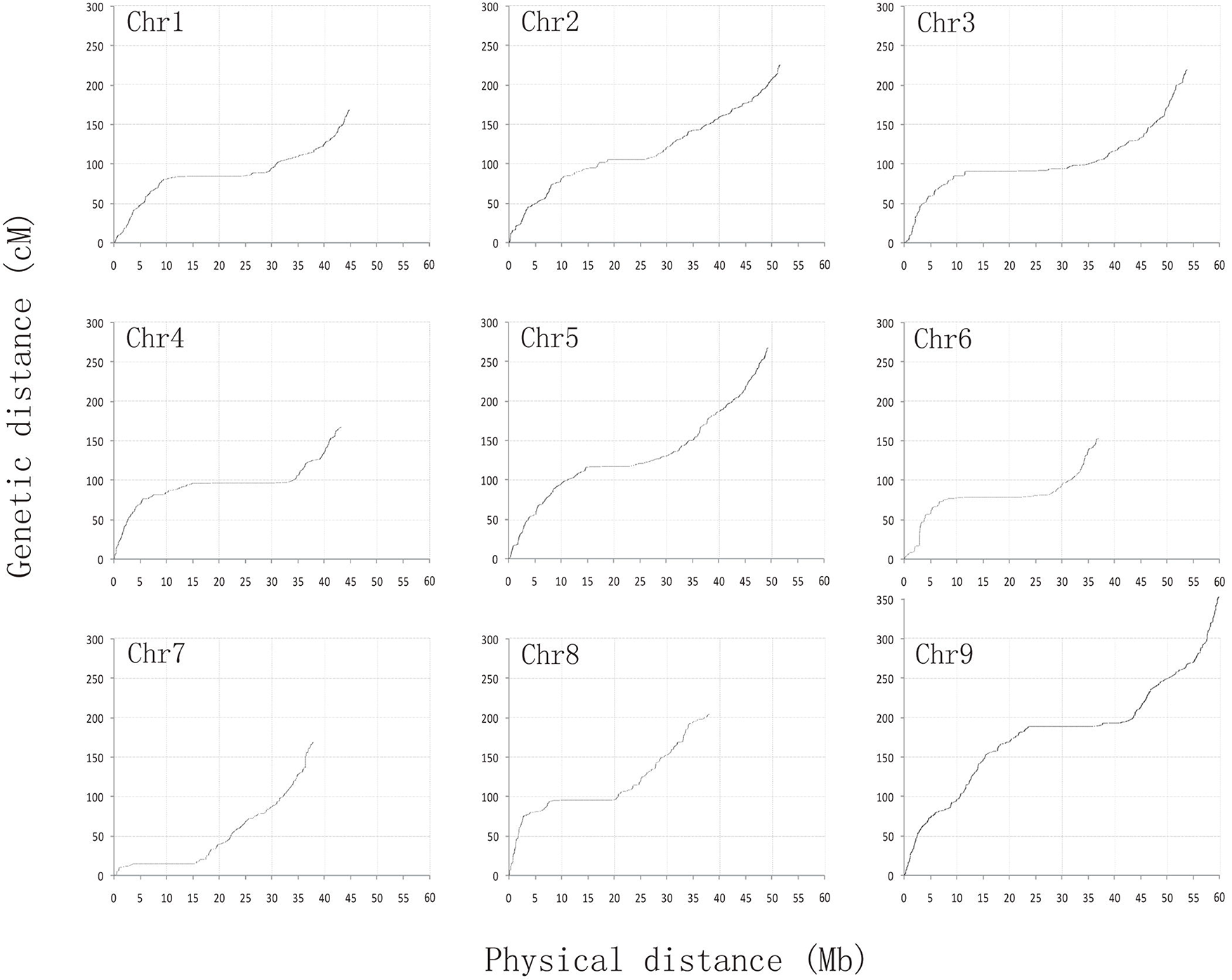
Genetic distance *vs.* physical distance. Genetic position of the 3437 bins is plotted against the corresponding physical position. Regions with low ratio of genetic distance to physical distance show heterochromatin regions

**Figure 3.**
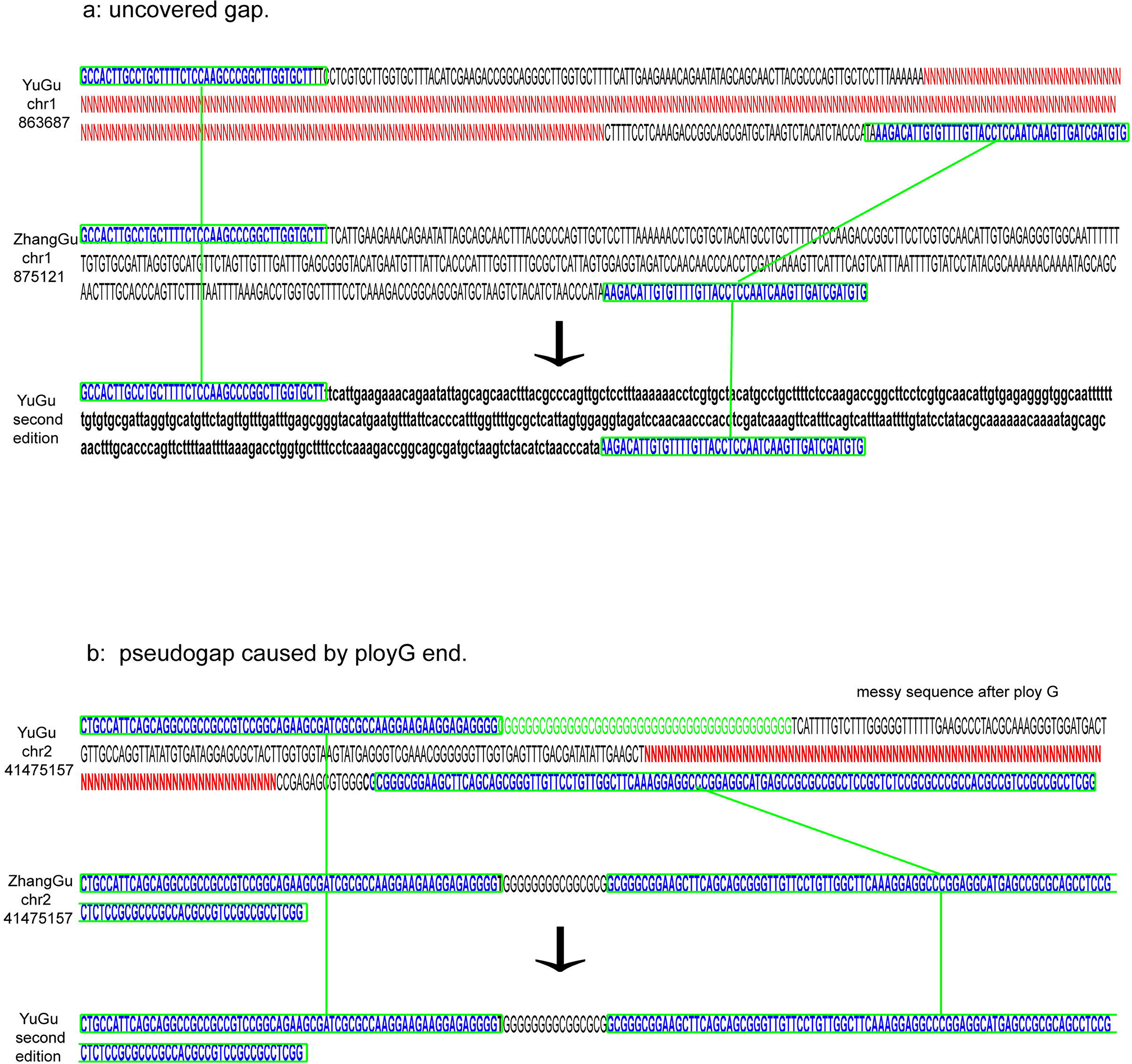
Gap filling in Yugu chromosome. Gap was filled by Zhanggu sequence when both flank sequence was match to Zhanggu chromosome. (**a**) Gap was caused by low coverage, both flank sequence was match to Zhanggu, gap was fill by lowercase Zhanggu sequence. (**b**) Gap was caused by ploy G sequence, Sanger sequencing can’t step over the ploy G.

**Figure 4.**
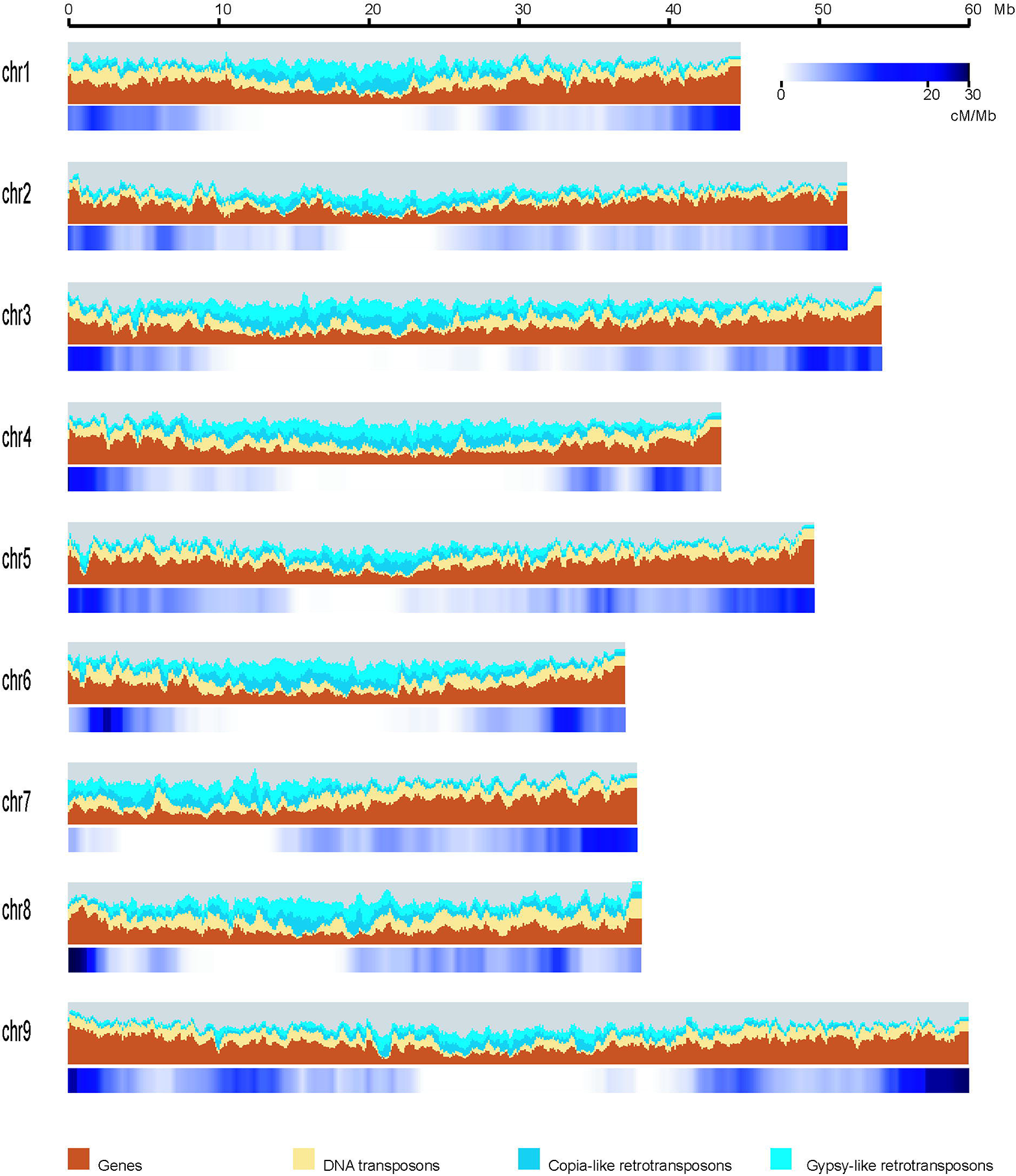
Genomic landscape of the Zhanggu chromosomes (second edition). Major DNA components are categorized into genes (brown), DNA transposons (yellow), Copia-like retrotransposons (dark blue), Gypsy-like retrotransposons (light blue), with respective DNA contents of 19%, 13%, 10% and 21% of the genome sequence. Categories were determined for 1-Mb windows with a 0.2-Mb shift. Recombination ratio was shown in blue bars, ranging from 0 cM/Mb to 30 cM/Mb.

**Table 2.**
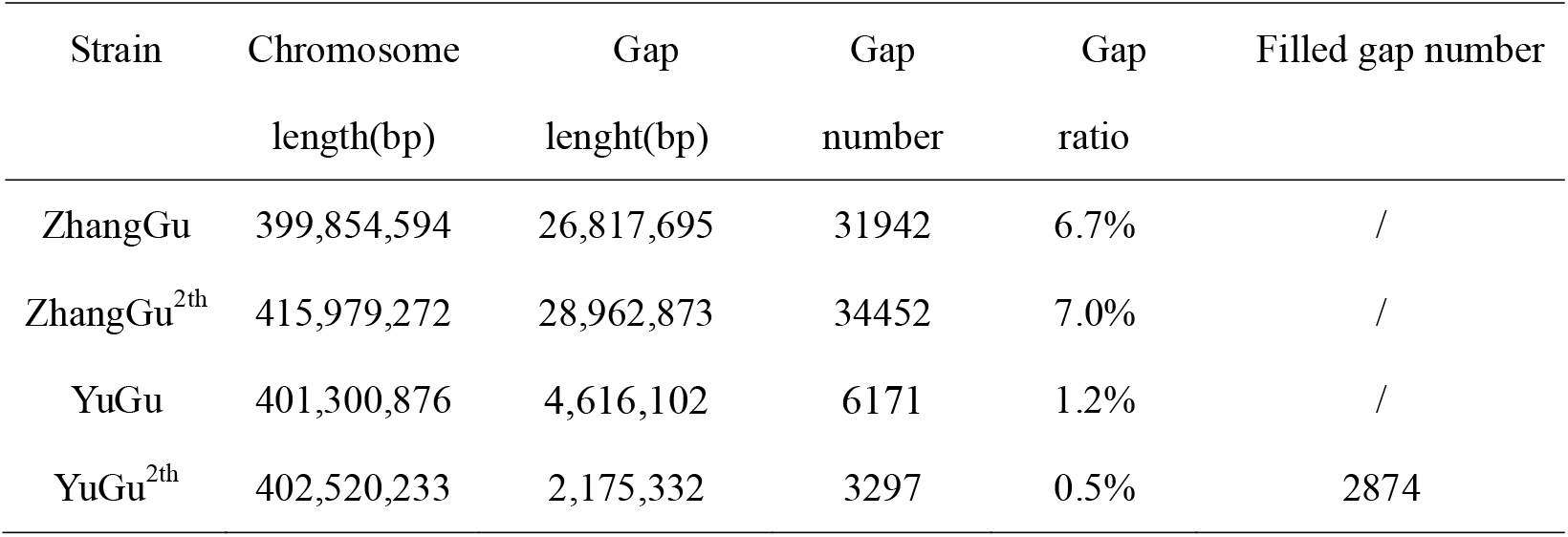
Summary of Zhanggu second edition and Yugu second edition

| Strain | Chromosome<br>length(bp) | Gap<br>length(bp) | Gap<br>number | Gap<br>ratio | Filled gap number |
| --- | --- | --- | --- | --- | --- |
| ZhangGu | 399,854,594 | 26,817,695 | 31942 | 6.7% | / |
| ZhangGu<sup>2th</sup> | 415,979,272 | 28,962,873 | 34452 | 7.0% | / |
| YuGu | 401,300,876 | 4,616,102 | 6171 | 1.2% | / |
| YuGu<sup>2th</sup> | 402,520,233 | 2,175,332 | 3297 | 0.5% | 2874 |

To identify agronomic traits related loci which were important in foxtail millet, gene mapping and QTL analysis was done based on this RIL population (WU *et al*. 2008; HIRANO *et al*. 2011). Nine agronomic traits, which can be divided into two categories: qualitative traits (Sethoxydim resistance, leaf color, bristle color, anther color, tassel hardness) and quantitative traits (plant height, heading date, flag leaf width, flag leaf length), **were measured with replications** (NACIRI *et al*. 1992).

All five qualitative traits show single gene control pattern (Fig. 5). Leaf color (green - yellow) was controlled by a dominant gene *Z3lc* mapped onto the long arm of chromosome 7 (bin2535); Sethoxydim resistance (resistance - sensitive) was controlled by a dominant gene *Z3sr* mapped onto the short arm of chromosome 7 (bin2346); bristle color (red - green) was controlled by a dominant gene *Z3bc* mapped onto the short arm of chromosome 4 (bin1436); anther color (yellow - brown) was controlled by a dominant gene *Z3ac* mapped onto the long arm of chromosome 6 (bin2304); tassel hardness (stiff - flexible) was controlled by a dominant gene *Z3th* mapped onto the short arm of chromosome 5 (bin2027).

**Figure 5.**
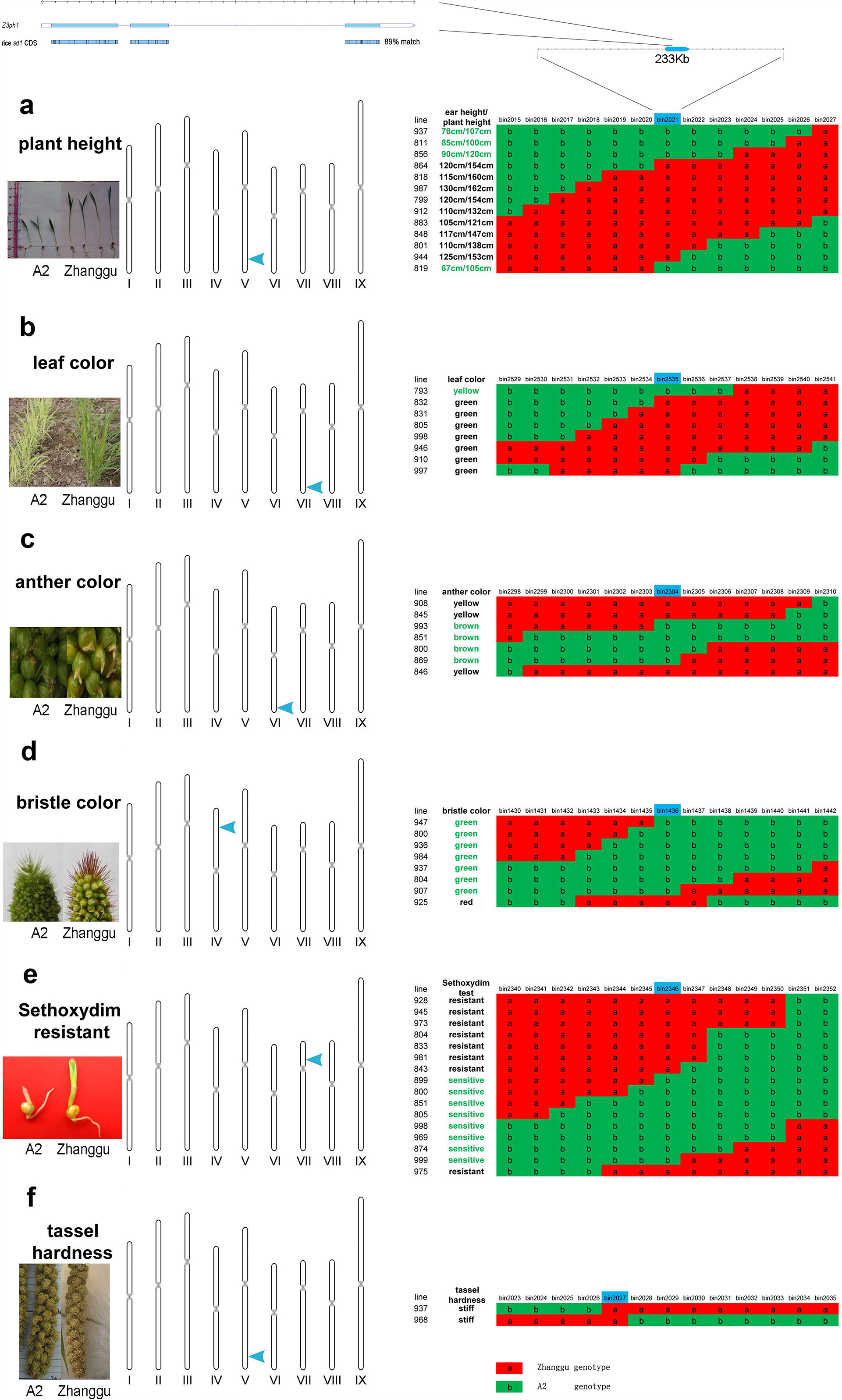
Gene mapping of the largest effect locus of plant height and five qualitative traits in foxtail millet. Genotype of recombination lines are shown in red and green block, “a” in red block means paternal genotype, “b” in green means maternal genotype. Phenotype of recombination lines are shown in the left of genotype blocks. (**a**) Gene mapping of the largest effect locus of plant height (paternal: tall – maternal: dwarf). (**b**) Gene mapping of the locus of leaf color (paternal: green – maternal: yellow). (**c**) Gene mapping of the locus of anther color (paternal: yellow – maternal: brown). (**d**) Gene mapping of the locus of bristle color (paternal: red – maternal: green). (**e**) Gene mapping of the locus of Sethoxydim resistance (paternal: resistant – maternal: sensitive). (**f**) Gene mapping of the locus of tassel hardness (paternal: stiff – maternal: flexible).

Using 184 RIL lines and a F2 population (the F2 population used to construct linkage map in 2009), we detected two quantitative trait loci (QTL) related to plant height (chr2, chr5, Fig. 6 b). The largest effect locus (25.3% in 2011; 8.8% in 2010; 46.3% in 2009) was then mapped onto the bin2021 (Fig. 5 a). The candidate gene *Z3ph1* in bin2021 shown 89% identity to the known rice gibberellin-synthesis gene *sd1* (SASAKI *et al*. 2002) (Fig. 5 a, Fig. 6 a), which indicated the plant height in millet may also control by GA20ox. We also detected three quantitative trait loci (QTL) related to heading date (chr2, chr7, chr9). One locus which is identical to the position of the leaf color control gene *Z3lc*, which indicated that the leaf color also affect the heading date ( Fig. 6 b). Flag leaf width and length are more complex than plant height and heading date. We detected five large effect QTLs related to flag leaf length, but no one show repeated emergence in three years. QTLs related to flag leaf width are tiny and irregular, but one QTL located in chromosome 9 repeated in three years.

**Figure 6.**
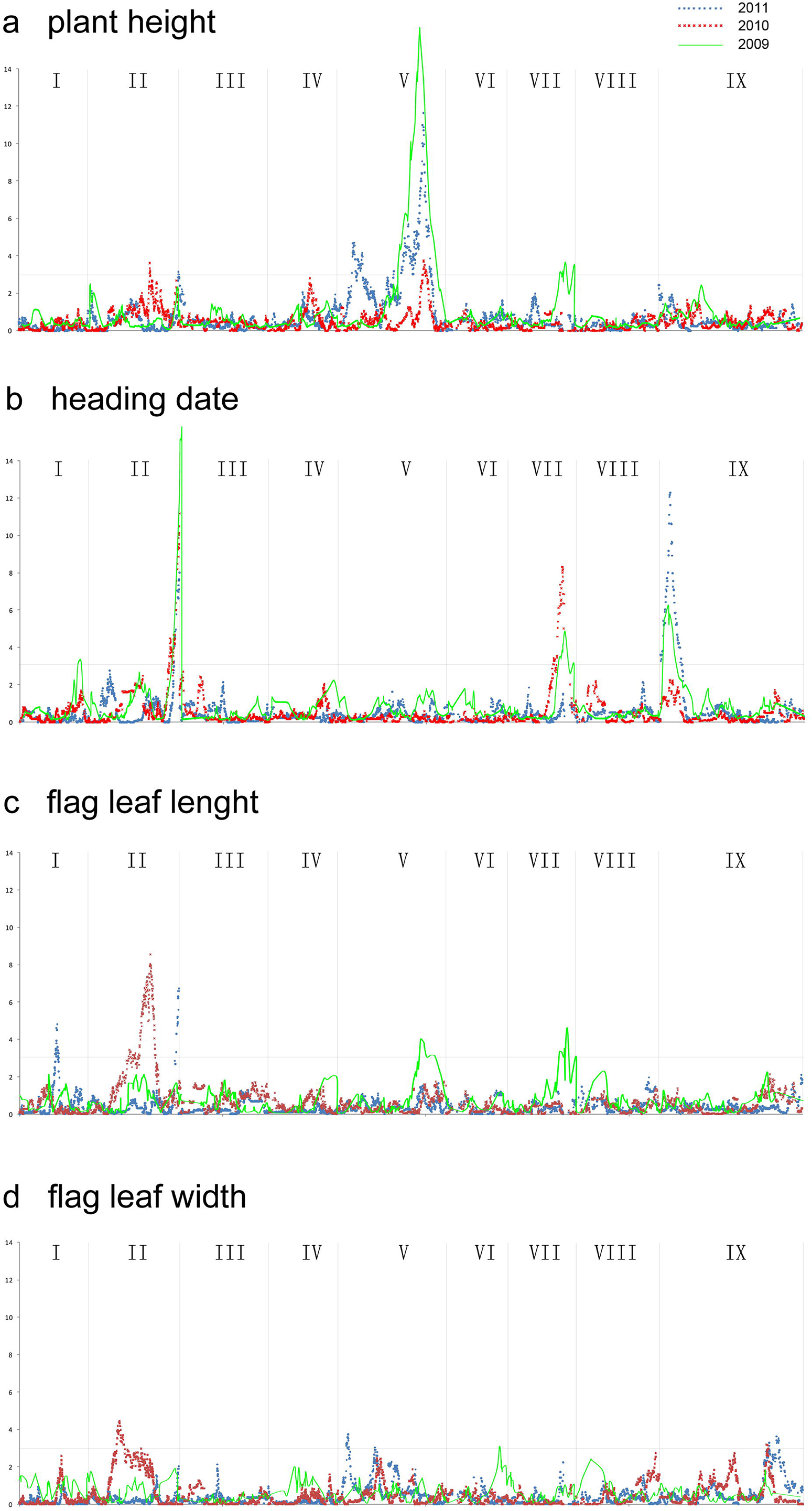
QTL analysis of five quantitative traits in foxtail millet. Peak signals of three years date are shown in three colors. (**a**) QTL analysis of plant height. (**b**) QTL analysis of heading date. (**c**) QTL analysis of flag leaf length. (**d**) QTL analysis of flag leaf width. Lines in green means data collected in 2009, lines in red means data collected in 2010, lines in blue means data collected in 2011.

## DISCUSSIONS

Grasses provide staple food for the vast majority of the world population. It can be divided into different tribes (DOUST *et al*. 2009; LATA *et al*. 2013). Foxtail millet belongs to one of the tribe which contains many drought-resistant species, such as switchgrass (*Panicum virgatum*), pearl millet (*Pennisetum glaucum*), prosomillet (*Panicum miliaceum*). Although foxtail millet was one of the most drought-tolerant crops, it was still substituted by high yield bybrid corn. The most important reason was due to the relative low productivity and high labor cost of the traditional foxtail millet. Data showed that the foxtail millet also had distinct heterosis between different individuals, which was similar with the other grass crop rice (SILES *et al*. 2004; HUANG *et al*. 2010). The high yield, herbicide resistance characteristics made hybrid millet suitable for large scale planting and industrilization (DOUST *et al*. 2004; JIA *et al*. 2007; WANG *et al*. 2012; JIA *et al*. 2012)..

In this paper, the genome draft map, high density genetic linkage map and QTL mapping of several important agronomic trait were done using next generation sequencing (ZHANG *et al*. 2012) and Sanger sequencing (BENNETZEN *et al*. 2012) method until 2012.

We developed the bin map and high density SNP markers, of which the data has been made publicly available. We also provided gene mapping and QTL mapping of nine important agronomic traits, the peak signals at nine loci will provide an important tool to foxtail millet breeding. What is more, we update the millet draft genome to the millet fine map by adding unseated scaffolds (16Mb data). The Zhanggu and Yugu genome reference updated in this work will be helpful for millet resequencing and GWAS analysis in future. Our study also indicates that resequencing of RILs could provide an effective approach for high quantity genome assembly and gene mapping.

## ACKNOWLEDGMENTS

We are grateful to all participants of the Agriculture department at BGI. The work was supported by the National Key Technology R&D Program (2015BAD02B01-7), the Technology Innovation Program Support by Shenzhen Municipal Government (JSGG20130918102805062 and CXZZ2015033017181006), the Basic Research Program Support by Shenzhen Municipal Government (JCYJ20150831201123287 and JCYJ20120618172523025). The funders had no role in study design, data collection and analysis, decision to publish, or preparation of the manuscript.

## Availability of supporting data

The genome sequence and annotation data set of Zhangu (second edition) has been deposited into NCBI (accession number: PRJNA73995). The genome sequence and annotation data set of Yugu (second edition) has been deposited into NCBI (accession number: PRJNA80183). The genome reference sequence and genotype of 184 RILs can be downloaded from ftp://ftp.genomics.org.cn/pub/Foxtail_millet (to be uploaded and cited from GigaDB if it passes review). The details of 3437 bins can be found in Table S2.

## Supporting information

**Table S1 MSG sequencing and Recombination events information (XLS)**

**Table S2 SNPs information generated from F2 population.**The document includes the genotypes of samples. The missing genotype is markers as “-” specially. Format description (left to right): Column1: Chromosome name.Column2: Position. Column3: Genotype of two parents and each F2 sample.

**(XLS)**

